# The influence of 4-thiouridine labeling on pre-mRNA splicing outcomes

**DOI:** 10.1101/2021.09.03.458914

**Authors:** Jessie Altieri, Klemens J. Hertel

## Abstract

Metabolic labeling is a widely used tool to investigate different aspects of pre-mRNA splicing and RNA turnover. The labeling technology takes advantage of native cellular machineries where a nucleotide analog is readily taken up and incorporated into nascent RNA. One such analog is 4-thiouridine (4sU). Previous studies demonstrated that the uptake of 4sU at elevated concentrations (>50µM) and extended exposure led to inhibition of rRNA synthesis and processing, presumably induced by changes in RNA secondary structure. Thus, it is possible that 4sU incorporation may also interfere with splicing efficiency. To test this hypothesis, we carried out splicing analyses of pre-mRNA substrates with varying levels of 4sU incorporation (0-100%). We demonstrate that increased incorporation of 4sU into pre-mRNAs decreased splicing efficiency. The overall impact of 4sU labeling on pre-mRNA splicing efficiency negatively correlates with the strength of splice site signals such as the 3’ and the 5’ splice sites. Introns with weaker splice sites are more affected by the presence of 4sU. We also show that transcription by T7 polymerase and pre-mRNA degradation kinetics were impacted at the highest levels of 4sU incorporation. Increased incorporation of 4sU caused elevated levels of abortive transcripts, and fully labeled pre-mRNA is more stable than its uridine-only counterpart. Cell culture experiments show that a small number of alternative splicing events were modestly, but statistically significantly influenced by metabolic labeling with 4sU at concentrations considered to be tolerable (40 µM). We conclude that at high 4sU incorporation rates small, but noticeable changes in pre-mRNA splicing can be detected when splice sites deviate from consensus. Given these potential 4sU artifacts, we suggest that appropriate controls for metabolic labeling experiments need to be included in future labeling experiments.

## Introduction

4-thiouridine (4sU), a uridine analog, is a commonly used metabolic label that can be employed to investigate a wide variety of topics that range from pre-mRNA generation to mRNA degradation. When added to cell medium, 4sU is readily taken up and incorporated into newly transcribed RNA [1,2]. In mammalian cells, this uptake results in a 0.5% to 2.3% median 4sU incorporation rate [3,4] when in the presence of 500µM 4sU for 2hr and 100µM for 24hr. 4sU differs from uridine by the existence of a sulfur atom rather than an oxygen atom in the 4^th^ carbon position on the nitrogenous base [5]. 4sU incorporation into RNA assists in the physical or computational isolation of RNA species. The modified nucleotide can be biotinylated to allow for bead capture and subsequent isolation [1,6]. The presence of 4sU in RNA also induces a single nucleotide sequencing artifact that can later be used to identify the presence of 4sU in an RNA strand [3,6]. Despite its widespread use, not much is known about the functional implications of 4sU incorporation in downstream RNA processes. Recent studies suggested that labeling at 4sU concentrations >100µM, concentrations commonly used across many different cell types[2], may elicit adverse effects, especially during extended labeling times (>12hrs) [7]. Indications of toxicity were also observed at concentrations as low as 50µM [7], presumably because ribosomal RNA synthesis is inhibited, triggering cellular stress response [7]. These observations suggested that 4sU incorporation can negatively impact translation and cellular homeostasis. Given the extensive use of 4sU labeling in the study of nascent pre-mRNA splicing, we set out to determine if intron removal efficiency is influenced by 4sU incorporation.

Pre-mRNA splicing is an essential biological process that allows for proteomic complexity, despite the relatively small number of protein-coding genes within the human genome [8]. This process is carried out by the spliceosome, a multiprotein/snRNA complex that recognizes and binds to hallmark pre-mRNA sequences. These sequences include both the 5’ and 3’ splice sites (ss), the branch point, and the polypyrimidine tract [9]. A single nucleotide change in any of these sequences can alter the efficiency of pre-mRNA recognition by the spliceosome. Thus, it is possible that 4sU incorporation impacts pre-mRNA splicing.

Using complementary *in vitro* and cell culture experiments, we show that pre-mRNA splicing can be impacted by the incorporation of 4sU into pre-mRNAs. The extent of splicing interference depends on 4sU incorporation levels and the intrinsic ability of introns to be recognized by the spliceosome. Thus, at high levels of 4sU incorporation, weaker exons are more likely to be differentially processed. 4sU labeling also interfered with *in vitro* transcription and *in vitro* pre-mRNA stability at elevated levels of incorporation. We conclude that the presence of 4sU in nascent RNA can interfere with pre-mRNA splicing. However, the negative effects are only detectable at elevated levels of 4sU incorporation levels and in the context of weak splicing events.

## Materials and Methods

### *In vitro* transcription of radiolabeled pre-mRNA

ADML and β-Globin, two well-studied DNA constructs that are spliced with ease by the spliceosome when transcribed under normal conditions, were used to generate pre-mRNA for *in vitro* splicing[10]. Splice site strengths for each of the constructs were determined using MaxEntScanner [11]. After linearizing with BamHI (**Promega, R6021**) and cleaning/concentrating (**Zymo, D4013**) ADML and β-Globin, a 10µL radioactive *in vitro* transcription reaction was performed using T7 polymerase, [α-^32^P] CTP (**Perkin Elmer, BLU008H250UC**), and phosphorylated 4-thiouridine (**TriLink, N-1025**). Different strands of pre-mRNA were generated for each construct that included different quantities of 4sU. The amounts of 4sU relative to the total amount of rUTP used in the transcription reaction mixture were 0%, 2.5%, 15%, 30%, and 100%. The resulting pre-mRNA was PAGE purified, eluted, precipitated, and resuspended at 10µL per 100,000cpm on the Geiger counter.

### *In vitro* pre-mRNA splicing

*In vitro* splicing was carried out by following a well-established protocol [12] in either 12.5µL or 25µL reactions for 90min or the indicated time for the experiment. The splicing reactions were run out on 6% polyacrylamide (19:1 acrylamide: bisacrylamide) 7 M urea gels and imaged using a GE Typhoon imager. Splicing efficiency was determined by following the “percent spliced” equation in the *in vitro* splicing protocol [12]. Splicing rates were determined by fitting data points to pseudo-first-order decay equations. All resulting gels were analyzed using GelAnalyzer 19.1 software [13], visualized using GraphPad Prism9 software [14] and all statistical analyses were performed with Prism9’s built-in statistical analysis tools. The degradation assays were performed identically to the *in vitro* splicing assays, except we created degradation conditions by omitting the addition of RNasin, CP, and ATP from the reaction mixture and by depleting the nuclear extract of ATP by incubation at room temperature for 30min prior to the addition of the labeled RNA. Normalized degradation profiles were calculated by dividing the band intensity of the intact pre-mRNA by the band intensity of the degraded pre-mRNA along the time points of the reaction.

### HEK293 metabolic labeling

HEK293 cells (**ATCC, CRL-1573**) were grown in 6-well plates at 37°C, 5% CO2 in a humidified incubator to 80% confluency for the start of the 2hr labeling experiments and 40% confluency for the start of the 24hr labeling experiments. The cells were grown in DMEM/High Glucose (**HyClone, SH30022.01**) plus 10% FBS (**GeneClone, 25-550**). A metabolic labeling protocol was followed[6] and the experimental HEK293 cells were incubated in the presence of 40µM unphosphorylated 4-thiouridine (**Cayman Chemical, CAY16373-5**). Total RNA was harvested from the cells using Trizol (**Ambion, 15596018**) and processed per the metabolic labeling protocol. The resulting pure total RNA was stored in TE at -80°C for further experimentation.

### Endpoint PCR

Using 1µg of total RNA harvested from the metabolic labeling experiments, cDNA was generated using a first-strand cDNA synthesis kit (**Invitrogen, 18080-044**) by following the manufacturer-provided protocol. Oligo d(T)16 (**IDT, 51-01-15-06**) was used as the primer for the cDNA reaction. PCR primers were designed that flank the exons/regions of interest or using previously described primer sets [15]. Primer sequences are as follows: **F_RIOK3_Ex5** CCGGTTCCCACTCCTAAAAAGGGC; **R_RIO-K3_Ex10** CCAGCATGCCACAGCATGTTATACTCAC; **F_TRA2B_Ex1** AGGAAGGTGCAAGAGG TTGG; **R_TRA2B_Ex3** TCCGTGAGCACTTCCACTTC; **F_CLK2_Ex3** CCGGACATTTAGCCGC TCAT; **R_CLK2_Ex6** TGGCCATGGTAGTCAAACCA; **F_DDX5_Ex11** ATTGCTACAGATGTGG CCTCC; **R_DDX5_Ex12** TGCCTGTTTTGGTACTGCGA. We designed additional sets of primers to investigate both alternative ‘Alt’ splicing events and constitutive ‘Const’ splicing events: **F_ADAP2_Alt_Ex2** CAGCAGAGTTAAATCTGTGCGAC; **R_ADAP2_Alt_Ex4** CTTAGCTCGAATCCATTGTTCC; **F_ADAP2_Const_Ex6** AATGCCACCTTCCAGACAGA; **R_ADAP2_Const_Ex8** TGGCCCAGTCTTTTCCATGA; **F_DOLPP1_Alt_Ex5** GCACCAAACAAACAACGCCA; **R_ DOLPP1_Alt_Ex6** GGAAGAACTCGGAGACAGGC; **F_ DOLPP1_Const_Ex2** GTCGGTTTCGTGACCCTCATC; **R_ DOLPP1_Const_Ex5** CAGGAACCTGGCGTTGTTTG; **F_ZNF711_Alt_Ex6** CACCAGTGGACATTCAGTAGC; **R_ ZNF711_Alt_Ex8** GCTTGACAATCTTCATACCTTCG; **F_ ZNF711_Const_Ex4** GTGATTCAAGCAGCTGGAGG; **R_ ZNF711_Const_Ex6** CATCTTCTCCCGCTGCATTC. PCR was performed after appropriate adjustments in annealing temperature and elongation time were made for each primer set using an Eppendorf Mastercycler Nexus thermal cycler. PCR reactions were run out on 2.0% agarose gels at 120V for 40min and imaged using a Bio-Rad gel doc imager.

## Results

### The impact of 4sU on *in vitro* splicing efficiencies

To measure the effect of 4sU incorporation on pre-mRNA splicing we carried out *in vitro* and cell culture pre-mRNA splicing analyses. For the *in vitro* approach, we evaluated the pre-mRNA splicing activities of two well-studied minigene constructs, ADML and β-globin, by generating and testing pre-mRNAs with varying amounts of 4sU incorporation (0%, 2.5%, 30%, and 100% 4sU incorporation). ADML and β-globin minigenes are gold standard *in vitro* splicing constructs that have been shown to splice efficiently in splicing extract. Both minigenes harbor strong splice sites, with ADML containing a 5’ splice site with a MaxEnt score (MES) [11] of 7.9 and a 3’ splice site MES of 12.5, and β-Globin containing a 5’ splice site MES of 8.1 and 3’ splice site MES of 9.5 (Fig 1A). Given its higher 3’ splice site score, the ADML minigene is considered to be more efficiently spliced than the β-globin minigene.

**Fig 1.**
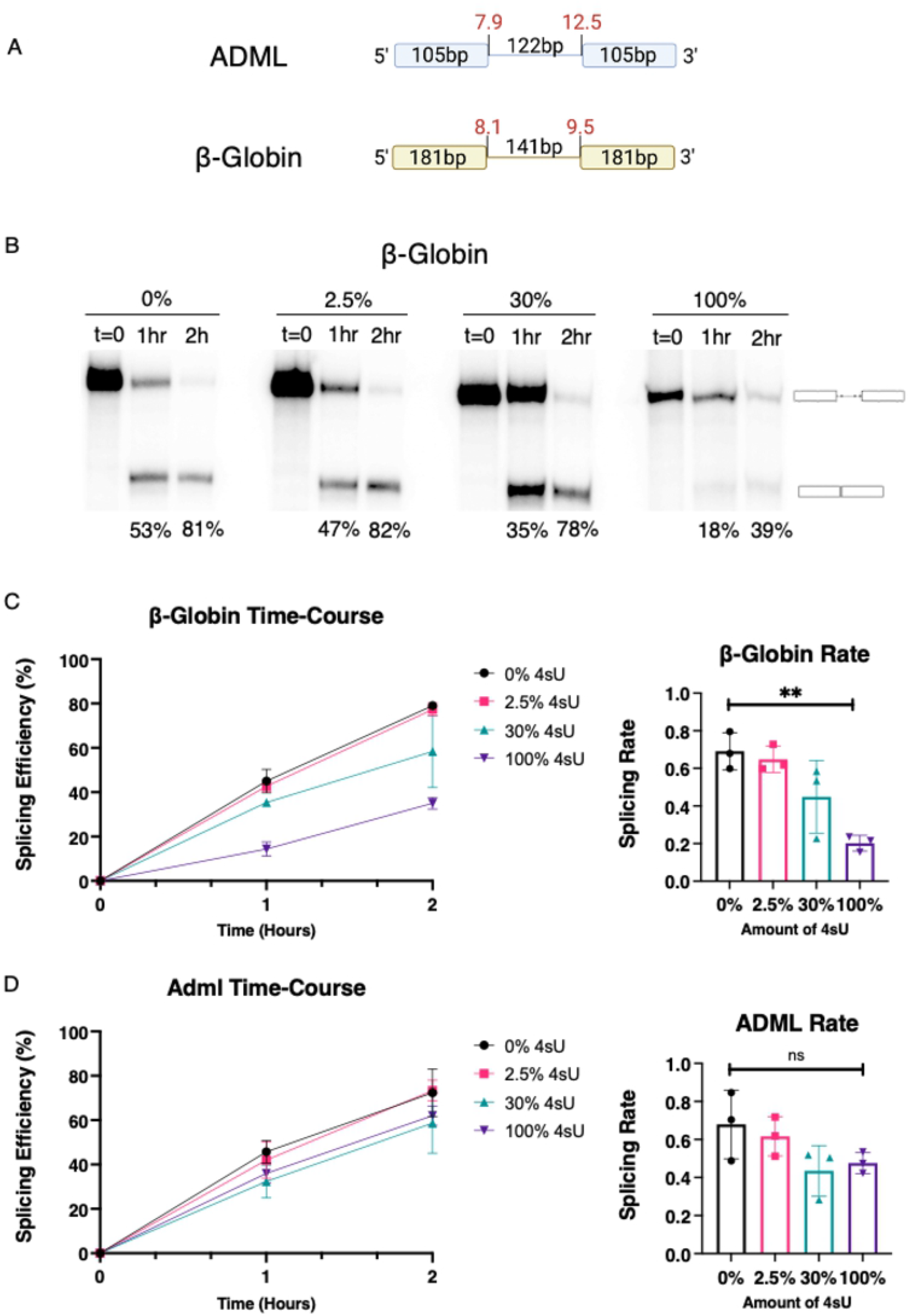
*In vitro* splicing kinetics of 4sU labeled RNA. (A) Schematic depictions of the ADML and β-Globin minigenes used. Exon and intron sizes and splice site strengths (MaxEnt score) are indicated. (B) Autoradiogram representing time course behavior of 0%, 2.5%, 30%, and 100% 4sU containing β-Globin pre-mRNAs. Precursor RNA and spliced products are defined on the right of the gel. The numbers under each lane represents the efficiency of splicing. (C) Graphical representation of β-Globin time-course splicing assay splicing efficiencies over time (n=3) and the corresponding splicing rate (ordinary one-way ANOVA, P=0.0031). (D) Graphical representation of ADML time-course splicing assay splicing efficiencies over time (n=3) and the corresponding splicing rate (ordinary one-way ANOVA, P=0.14)

To determine if 4sU incorporation influences rates of *in vitro* splicing, splicing efficiencies after 1hr and 2hrs were compared between minigene constructs with incorporation rates of 0%, 2.5%, 30%, and 100% 4sU. For β-globin we observe a gradual decrease in the average splicing efficiency as the level of 4sU incorporation in the pre-mRNA increases (Figs 1B and C). While 4sU incorporation rates of 2.5% did not change observed splicing rates significantly, higher levels of 4sU incorporation led to an up to a 3-fold reduction in splicing (Fig 1C). The observations are qualitatively similar for the tested ADML pre-mRNA (Fig 1D). However, considering the range of splicing reduction and variation of the independent experiments, the differences observed did not meet the cutoff for statistical significance.

It is likely that the more striking differences in splicing efficiencies observed for the β-globin analysis are the consequence of its intrinsically weaker splice site signals when compared to ADML. We conclude that elevated levels of 4sU incorporation into pre-mRNAs can reduce splicing efficiencies and the resulting splicing rates, at least within the context of the time-course *in vitro* splicing assays used.

### Stability of 4sU Containing pre-mRNA

Modified RNAs are known to be more stable than unmodified RNAs [16,17]. To evaluate whether the incorporation of 4sU into RNA alters the stability of the resulting RNA we carried out *in vitro* degradation assays comparing the fate of fully modified or unmodified RNA samples over a 90 min period. Unmodified ADML RNAs displayed measurable degradation over the 90 min incubation (Fig 2A). Interestingly, fully modified ADML RNAs did not undergo any measurable degradation over the same time frame. Linear regression of the profiles indicates statistically significant differences between the degradation behaviors (Fig 2B, simple linear regression, P < 0.05). Analysis revealed a 9-fold difference in degradation kinetics between unmodified and fully modified RNA. These results suggest that the presence of 4sU promotes RNA stability, similar to other RNA modifications [18]. Just as 2’-O-methyl RNA and phosphorothioate RNA modifications resist nuclease activity [19], 4sU has the potential to limit nuclease attack.

**Fig 2.**
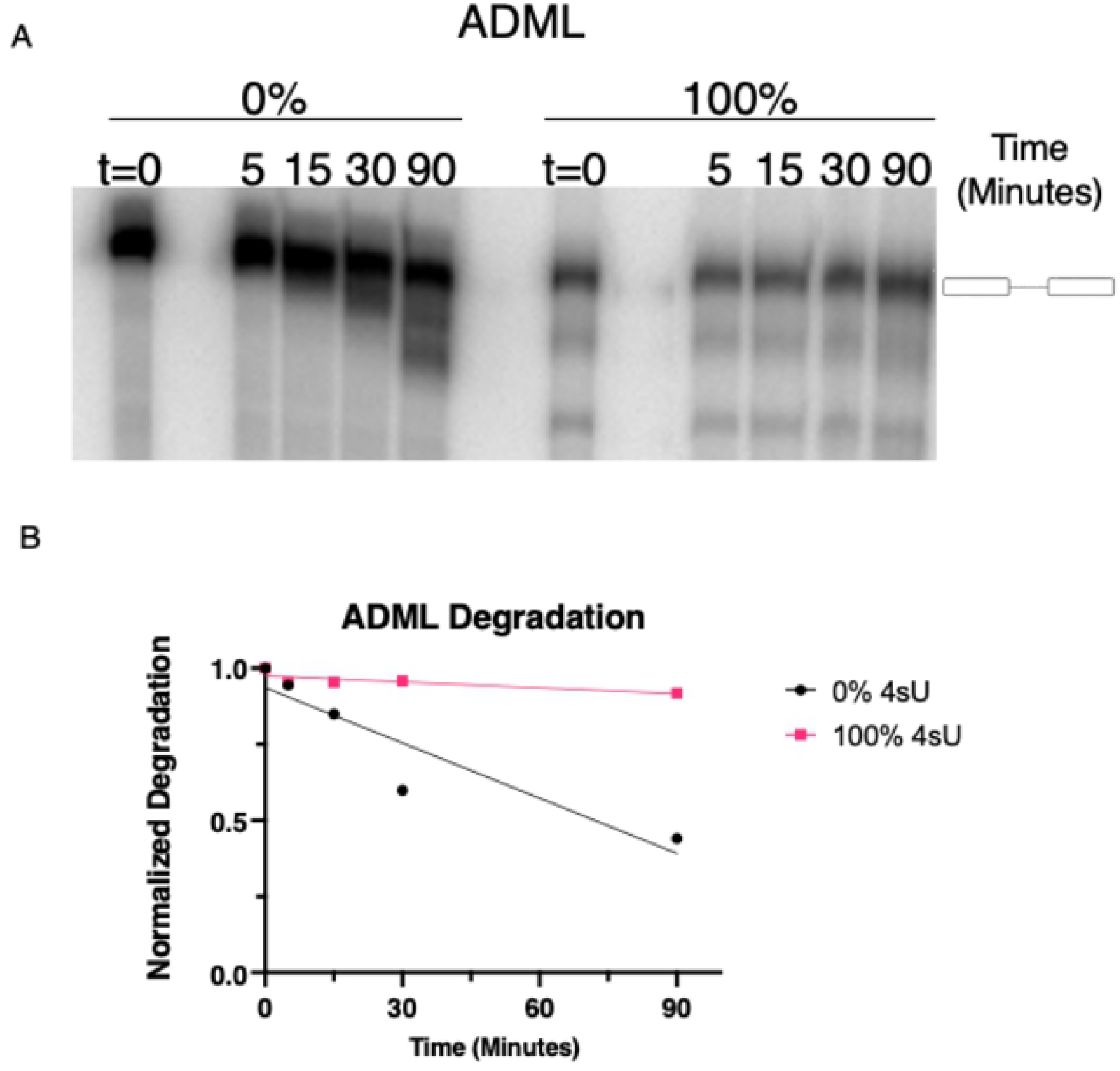
*In vitro* degradation assay of ADML minigene. (A) Representative autoradiograph of degradation time-course for unmodified and fully modified 4sU ADML pre-mRNA. t=0 represents input RNA. Full-length input RNA is defined by the cartoon to the right of the gel. (B) Quantitation of the data shown in (A). Simple linear regression of time course data was used for statistical analysis, with unmodified RNA P=0.024 and fully modified RNA P=.0092.

### The effects of 4sU on in vitro transcription

*In vitro* transcription of ADML and β-globin at the defined concentrations of 4sU resulted in variable accumulation of abortive transcripts. For example, transcription of ADML lead to a marked increase of an abortive transcript with higher incorporation rates of 4sU (Fig 3A). At 100% 4sU incorporation, the majority of transcripts were abortive, resulting in a truncated RNA.

**Fig 3.**
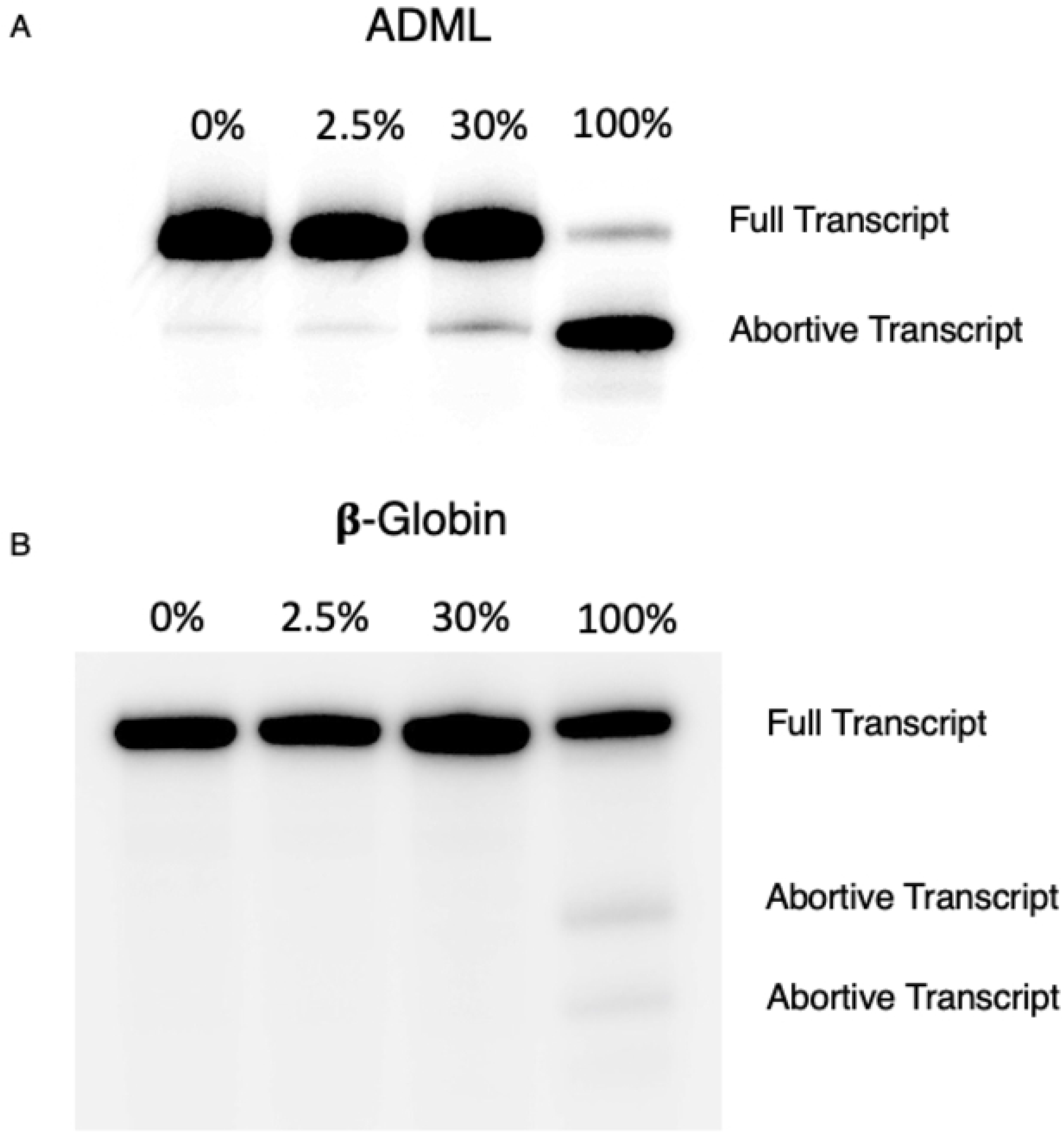
*In vitro* transcription of ADML and β-Globin. Autoradiograph depicting the T7 polymerase transcription profiles of A) ADML and (B) β-Globin. Four transcripts of each minigene were created with varying amounts, 0%, 2.5%, 30%, and 100%, of 4sU incorporated. The full pre-mRNA strands and abortive transcripts in each lane are labeled.

Qualitatively similar trends were observed when transcribing β-globin as multiple abortive transcripts are observed in the 100% 4sU lane (Fig 3B). Apparently, T7 RNA polymerase faced similar, but less extreme difficulties in generating the full β-globin pre-mRNA. These observations demonstrated that 4sU can negatively impact *in vitro* transcription processivity leading to pre-maturely terminated RNA products.

### The impact of 4sU on splicing in Hek293 Cells

To investigate the impact of 4sU incorporation on the splicing of endogenous genes we metabolically labeled HEK293 cells. Based on previous reports we used a 4sU concentration (40µM) that does not elicit toxic translation effects and we evaluated splicing after 2hrs and 24hrs of labeling. These 4sU metabolic labeling conditions are representative of the experimental approaches taken when evaluating nascent pre-mRNA splicing. They are also known to result in a 4sU incorporation rate of 0.5% to 2.3% [3,4]. In our analysis, we focused on evaluating the effects of 4sU on the splicing efficiency of constitutively spliced exons (considered efficient splicing events) and alternatively spliced exons (considered less efficient splicing events).

Our results demonstrate that 4sU metabolic labeling does not significantly interfere with the splicing of constitutive exons. For all examples tested we observed efficient exon inclusion regardless of the presence or absence of 4sU in either the 2hr or 24hr time points (Fig 4A). These observations suggest that efficient splicing events are not influenced by 4sU at incorporation rates typically used in cell culture experiments. To determine whether less efficient splicing events are affected by 4sU labeling we tested a select number of exon skipping events, alternative splice site selection events, and intron retention events. Most of the exon skipping events evaluated did not reveal any significant changes in splicing patterns between the control and 4sU labeled samples in either the 2hr or the 24hr samples (Fig 4B), even though some exon inclusion levels were intrinsically low (Fig 4B, ZNF711). As the only exception we detected a small, but statistically significant decrease in *DOLPP1* exon 6 inclusion (Fig 4B). We also detected small, but statistically significant changes in intron retention or alternative 5’ splice site choice after 24hrs of 4sU incubation (Figs 5A, B). At longer exposure to 4sU labeling, the level of *DDX5* intron 11 retention increased from 8% to 12%. While the magnitude of these changes is small, the increase in intron retention indicates that at the conditions used 4sU labeling can have an inhibitory effect on intron excision. Similarly, minor changes in alternative splice site choice are observed for *RIOK3* exon 8 after 24hrs of incubation (Fig 5B).

**Fig 4.**
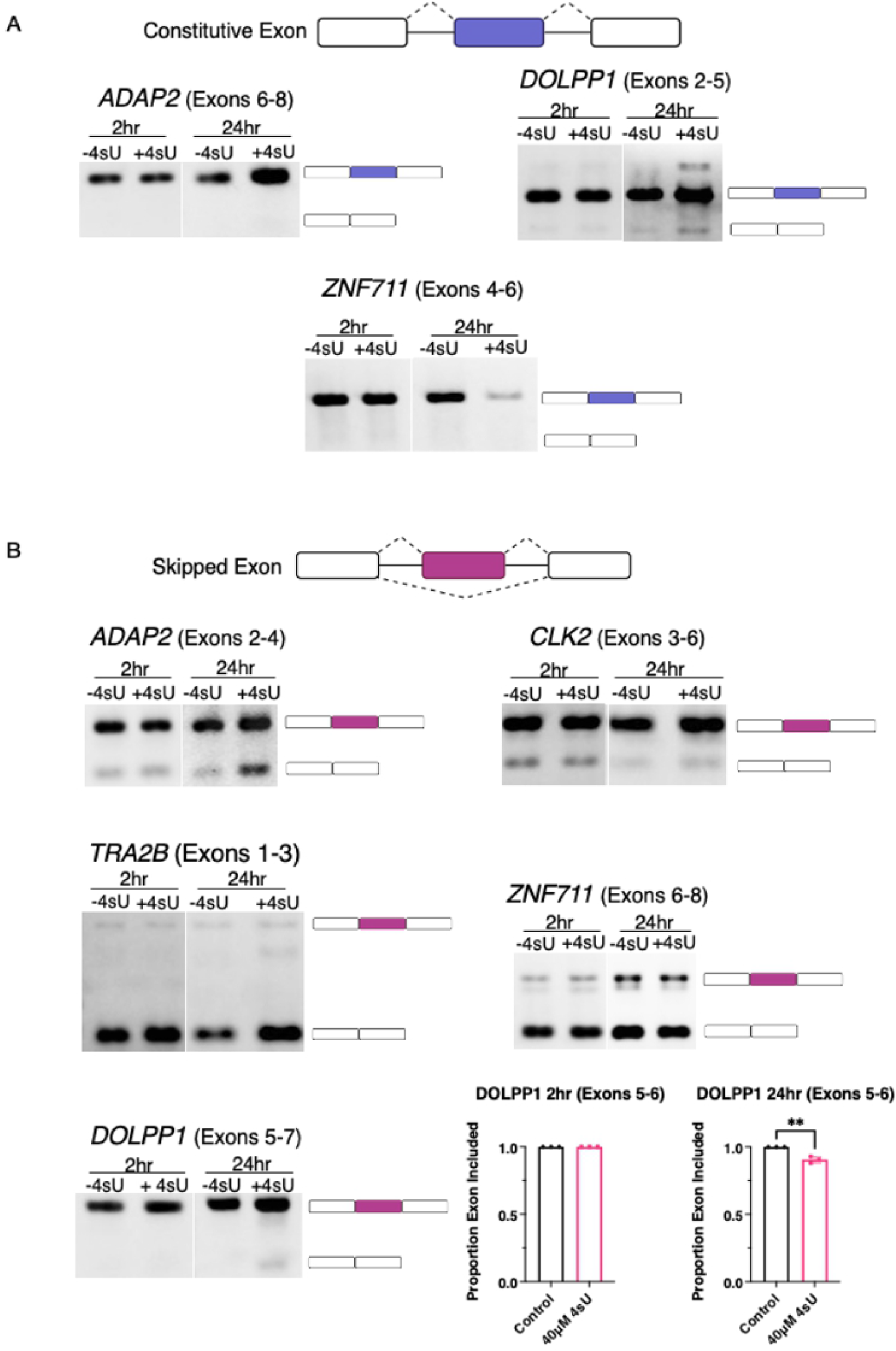
Splicing behavior in cell culture in the presence of 4sU. PCR analysis of (A) constitutive exons within the genes *ADAP2, DOLPP1*, and *ZNF711* and (B) alternatively included exons in genes *ADAP2, CLK2, TRA2B, ZNF711*, and *DOLPP1*. Alternative exon inclusion in *DOLPP1* (24hr sample t-test, P=0.0024). The cartoon to the right of each image indicates exon inclusion or exclusion. cDNA samples were analyzed after 2 and 24 hours of 4sU incubation.

**Fig 5.**
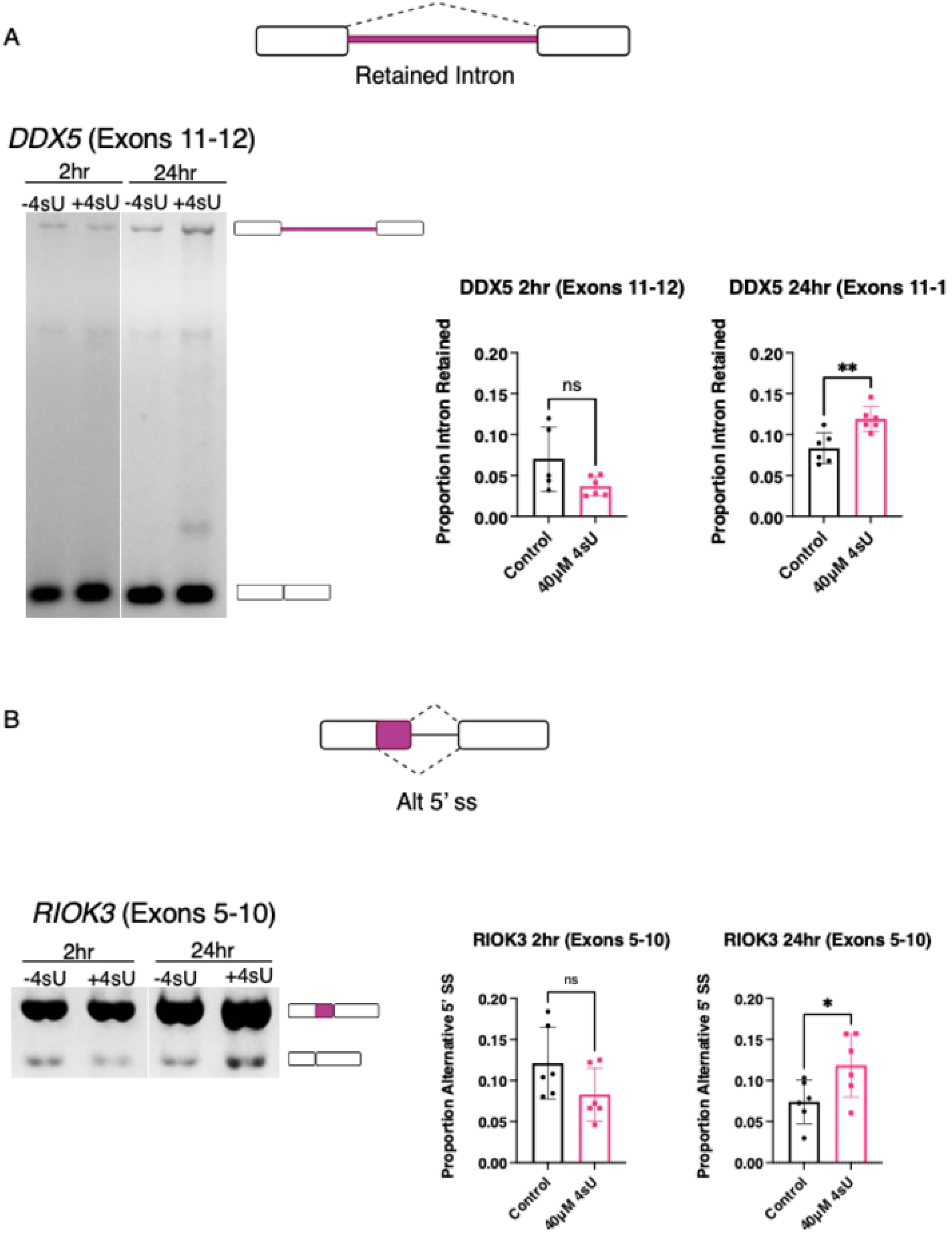
Alternative splicing behavior in cell culture in the presence of 4sU. PCR analysis of (A) intron retention in *DDX5* (2hr sample t-test, P=0.08; 24hr sample t-test, P=0.0045) and (B) alternative 5’ss selection in *RIOK3* (2hr sample t-test, P=0.012; 24hr sample t-test, P=0.042). The cartoon to the right of each image defines the alternative splicing products. cDNA samples were analyzed after 2 and 24 hours of 4sU incubation.

Consistent with our *in vitro* 4sU data we conclude that efficiently spliced introns are not significantly affected by the presence of 4sU, at least not at the conditions used. With a few exceptions, the alternative splicing events evaluated did not indicate altered splicing outcomes in the presence of 4sU. Thus, 4sU labeling at the 40 µM conditions used herein does not trigger drastic, widespread, and general changes in alternative splicing.

## Discussion

It is important to investigate whether the tools the scientific community use unintentionally create biases and side effects on the experiments being performed. 4sU labeling is a widely used tool in the RNA biology field to capture the processing of nascent RNAs and to evaluate RNA half-lives that is thought to have no consequence on the experimental outcome, aside from the previously reported inhibition of tRNA synthesis and nucleolar stress [7] at elevated concentrations. Here, we show that the incorporation of 4sU into pre-mRNA strands is not simply inert to the metabolism of an expressed RNA. At elevated levels of 4sU incorporation RNA transcription, pre-mRNA splicing, and RNA stability can be altered. However, at labeling concentrations typically used in cell culture experiments, the inhibitory effects of 4sU incorporation on pre-mRNA splicing are minimal, affecting only introns that are already weakly recognized by the spliceosome. Collectively, this data demonstrates that 4sU is not completely inert when employed as an experimental tool.

Interestingly, at high levels of 4sU incorporation transcription by T7 RNA polymerase resulted in increased levels of abortive transcripts. This was particularly striking for the ADML minigene (Fig 3A). To identify potential causes for the marked transcription interference we evaluated the RNA sequence of each transcript. The ADML minigene encodes a run of eight consecutive uridines towards the 3’ end of the intron. Thus, it is possible that when the polymerase encounters a run of consecutive uridines, the repeated incorporation of 4sU could interfere with polymerase processivity, resulting in increased abortive transcripts. The expected size of the ADML abortive transcript is consistent with this possible explanation. The longest stretch of consecutive uridines encoded by the β-globin minigene is only six nucleotides long. These observations suggest that longer runs of consecutive uridine induce abortive transcription when the RNA is labeled exclusively with 4sU. The potential implications of this transcription interference are straightforward. Incomplete pre-mRNA strands may not be stable, they may not be fully transcribed, and if they are, they could not have proper function.

The results of our experiments demonstrate that 4sU incorporation can affect pre-mRNA splicing. To what degree do these observations imply that the use of 4sU in metabolic labeling experiments should be viewed critically? We suggest considering the following aspects when designing metabolic labeling experiments. First, the concentration of 4sU used in the labeling approach. Second, the length of metabolic labeling, and third, the splice potential of the exons and introns investigated.

To evaluate the influence of 4sU labeling on pre-mRNA splicing we designed experiments that directly measure altered splicing outcomes using *in vitro* splicing assays. These investigations were complemented with 4sU labeling approaches of cell cultures, followed by mRNA analysis. Differences in intron excision, exon skipping, or alternative splice site selection rates in cell culture experiments can certainly be triggered by direct splicing effects, however, they can also be representative of altered mRNA stabilities and/or transcription outputs. With these interpretation limitations in mind, our data provide clear guidance when designing and interpreting splicing analyses using 4sU labeling. Increased levels of 4sU incorporation can lead to measurable defects in pre-mRNA splicing, preferentially for weaker splicing events. This is demonstrated in our *in vitro* splicing assays when pre-mRNAs of different incorporation levels were evaluated. However, our data also demonstrate that for the minigenes used, 4sU incorporation levels >30% need to be achieved to interfere with splicing efficiency. Such elevated levels of 4sU incorporation cannot be reached when performing cell culture metabolic labeling experiments. The length of 4sU incubation time is also an important factor when carrying out cell culture metabolic labeling. This is because longer incubation increases the fraction of mRNAs labeled with 4sU, allowing the establishment of steady-state levels of 4sU mRNAs. An important consideration when analyzing 4sU labeling experiments is the identity of the exon or intron evaluated. By definition, constitutive exons are always included in the final mRNA while alternative exons are not. As such, perturbations in recognizing exons, such as the presence of 4sU within the transcript, could have varying influences on alternative and constitutive exon inclusion levels. We observed that constitutive splicing is not affected by the presence of 4sU at the 40 µM concentration used. Most of the alternatively spliced exons evaluated were also not significantly affected by 4sU incorporation. Only a limited number of splicing events appeared to indicate altered splice site selection or reduced intron excision. Furthermore, the magnitudes of the observed splice changes were small, suggesting that while statistically significant, biological consequences are expected to be minimal. In summary, 4sU metabolic labeling of cell cultures has the potential to induce altered splicing. While infrequent, at small magnitudes, and only in the context of inefficiently spliced events, appropriate control experiments need to be included to deconvolute splicing differences of tested experimental conditions from those elicited by 4sU labeling.

It is known that naturally occurring 4-thiouridine modifications increase the stability of some organismal tRNAs[20] and it has been suggested that the incorporation of 4sU changes the structure of the transcribed RNA. The potential to form secondary structures has been shown to greatly impact exon recognition by the spliceosome [21], so it is entirely possible that 4sU incorporation into RNAs interferes with efficient exon recognition through inducing altered RNA structures.

The results presented here are not damning evidence against the use of 4sU, rather they are akin to a cautionary statement. 4sU does not broadly impact all aspects of pre-mRNA generation and processing, but it does impact those events that are more easily susceptible to change. If 4sU is to be used in an experiment, it is important to be mindful of the pre-mRNA splicing events that are being analyzed because a small number of event changes may be caused by the incorporation of 4sU itself.

## Acknowledgements

We are grateful to the members of our laboratory for helpful discussions and comments on this manuscript. Figures were in part created with Biorender.com.

